# Measuring Protein Insertion Areas in Lipid Monolayers by Fluorescence Correlation Spectroscopy

**DOI:** 10.1101/2020.09.23.310425

**Authors:** Jan Auerswald, Jan Ebenhan, Christian Schwieger, Andrea Scrima, Annette Meister, Kirsten Bacia

## Abstract

The insertion of protein domains into membranes is an important step in many membrane remodeling processes, for example in vesicular transport. The membrane area taken up by the protein insertion influences the protein binding affinity as well as the mechanical stress induced in the membrane and thereby its curvature. Total area changes in lipid monolayers can be measured on a Langmuir film balance. Finding the area per inserted protein however proves challenging for two reasons: The number of inserted proteins must be determined without disturbing the binding equilibrium and the change in the film area can be very small. Here we address both issues using Fluorescence Correlation Spectroscopy (FCS): Firstly, by labeling a fraction of the protein molecules fluorescently and performing FCS experiments directly on the monolayer, the number of inserted proteins is determined *in situ* without having to rely on invasive techniques, such as collecting the monolayer by aspiration. Secondly, by using another FCS color channel and adding a small fraction of fluorescent lipids, the reduction in fluorescent lipid density accompanying protein insertion can be monitored to determine the total area increase. Here, we use this method to determine the insertion area per molecule of Sar1, a protein of the COPII complex, which is involved in transport vesicle formation, in a lipid monolayer. Sar1 has an N-terminal amphipathic helix, which is responsible for membrane binding and curvature generation. An insertion area of (3.4 ± 0.8) nm^2^ was obtained for Sar1 in monolayers from a lipid mixture typically used in reconstitution, in good agreement with the expected insertion area of the Sar1 amphipathic helix. By using the two-color approach, determining insertion areas relies only on local fluorescence measurements. No macroscopic area measurements are needed, giving the method the potential to be applied also to laterally heterogeneous monolayers and bilayers.

**Statement of Significance:** We show that two color Fluorescence Correlation Spectroscopy (FCS) measurements can be applied to the binding of a protein to a lipid monolayer on a Langmuir film balance in order to determine the protein insertion area. One labelling color was used to determine the number of bound proteins and the other one to monitor the area expansion of the lipid monolayer upon protein binding. A strategy for the FCS data analysis is provided, which includes focal area calibration by raster image correlation spectroscopy and a framework for applying z-scan FCS and including free protein in the aqueous subphase. This approach allows determining an area occupied by a protein in a quasi-planar model membrane from a local, non-invasive, optical measurement.

## Introduction

Induction, maintenance and sensing of membrane curvature are essential processes for establishing the morphology of the cell and its organelles, as well as for shaping vesicles involved in cellular trafficking. Mechanisms to remodel the lipid bilayer into curved membranes include enzyme-mediated variations of the lipid composition, clustering of transmembrane proteins, the recruitment of scaffolding proteins and reversible insertions of hydrophobic protein motifs (1). The latter mechanism plays an important role in intracellular trafficking, where, among others, epsin and amphiphysin, components of the clathrin-mediated endocytic machinery as well as Arf1 and Sar1, which are small guanine nucleotide*-*binding proteins within the coat protein complexes COPI and COPII, utilize amphipathic helices to mediate membrane remodeling (2). These amphipathic helices are secondary structural motifs, which on one face harbor hydrophilic residues, while the opposite face is composed of hydrophobic residues that insert into the proximal leaflet of the membrane. The insertion of the amphipathic helix induces a high membrane curvature required for the formation of transport vesicles (3, 4).

COPII assembles on the endoplasmic reticulum membrane and forms vesicles responsible for transporting cargo from the ER towards the Golgi apparatus in the secretory pathway. As the amphipathic α-helix of Sar1 inserts only into the proximal leaflet of a membrane, phospholipid monolayers are useful model systems for analyzing this type of protein-lipid interactions in biological lipid bilayer membranes. Here we study the Sar1 protein (Sar1p) from baker’s yeast (*Saccharomyces cerevisiae*) in its permanently activated form with the non-hydrolyzable GTP analog GMP-PNP.

The mode of insertion of a particular protein or protein motif into the membrane, its insertion area and depth and orientation, constitutes vital information in the study of membrane remodeling. While a vast amount of literature exists regarding the insertion depth, fewer reports specifically deal with insertion areas (5-9). However, the lipid area the protein dislodges upon insertion provides complementary information regarding the protein-membrane interaction. It is related to the curvature the insertion will generate or the bending stress it can sustain. In addition, the insertion area can provide information towards the recognition of structural motifs that penetrate into the membrane apart from the actual hydrophobic motif (6, 9). Determining the area of insertion into membranes of varying composition aids in understanding protein-lipid interactions and binding affinity (5, 8).

Observing the interaction of proteins with lipid monolayers by means of a change in surface pressure or in surface area (10-12) constitutes a typical application of a Langmuir film balance. However, the determination of protein concentrations and diffusion properties is not possible on its own: Studying the microscopic properties of a system, such as insertion areas, usually requires transferring the film. For example, Huang *et al*. transferred the film to an ATR crystal to measure insertion areas (7). In doing so, the binding equilibrium is disturbed. Calculating insertion areas requires accurate knowledge of the amount of protein in the monolayer, preferably as a function of time and lateral position, to monitor the insertion process with respect to the completion and homogeneity, respectively.

To overcome these limitations inherent in the film balance setup, we combined the film balance with a confocal fluorescence correlation spectroscopy setup for optical detection of fluorescently labeled probes (Figure 1). Such a combined setup, pioneered by Gudmand *et al*. (13), allows FCS measurements in a monolayer as well as in the subphase and therefore the accurate quantification of the concentration of multiple distinct species *in situ*. Previously, FCS measurements were performed in solid-supported monolayers (14) or monolayers at liquid-liquid-interfaces with very small areas (15). Diffusion processes in monolayers were previously studied by SPT (single particle tracking) (16, 17), FRAP (fluorescence recovery after photobleaching) (18) and by radiochemical methods (19, 20). More recently, small troughs with a fixed area were used to study the diffusive behavior of surface-active molecules (21) or proteins adsorbed at a monolayer (22, 23).

**Figure 1:**
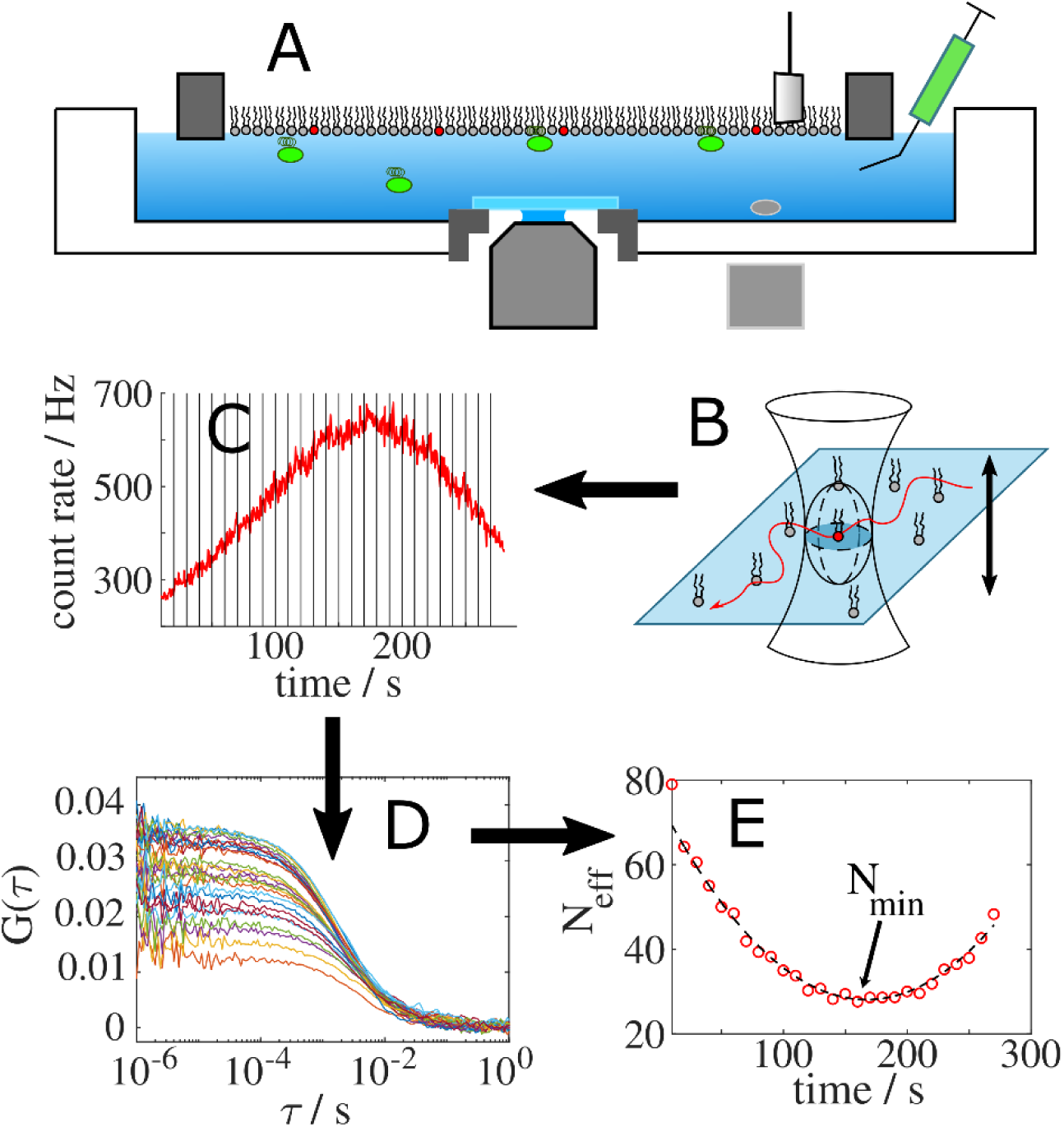
Schematic overview of the experimental setup and workflow to determine the number of fluorophores in the focus. (A) Side view of the film balance atop the microscope objective. Shown are the barriers on the left and the right of the lipid film (fluorescently labeled lipids in red), the Wilhelmy plate, the magnetic stirrer in the right compartment of the subphase (where the protein is being injected under the barrier from a syringe) and the free and bound Sar1-molecules in the subphase (depicted in green). (B) Fluorescent lipids diffusing in the membrane plane through the focus. The outer hyperboloid depicts the focused laser beam and the inner ellipsoid depicts the effective focus relevant for FCS measurements. The monolayer slowly passes through the focus as the subphase water evaporates. FCS measurements of labeled protein and labeled lipids can be acquired. Here, a fraction of protein was labeled in green and, where indicated, a fraction of lipid in red. (C) The count rate trace recorded during one focal passage of the film is split into smaller time windows for analysis. (D) Correlating the fluorescence fluctuations in each time window results in a number of autocorrelation curves that are fitted to a model to extract the particle number and diffusion times. (E) The fitted particle numbers along the focal passage follow a parabola with the true particle number at the minimum.

Fluorescence Correlation Spectroscopy (FCS) was first introduced in the 1970s and is a low-invasive, single-molecule sensitive fluctuation technique. Today it is a versatile and robust technique that is widely applicable to biologically relevant macromolecules such as proteins and nucleic acids as well as lipid assemblies. Because fluorescent labels can be specifically coupled to individual molecules, fluorescence fluctuation techniques are highly specific and sensitive and permit analysis of interactions even in complex systems including membrane preparations and living cells. Advantages in the context of monolayer measurements include the high spatial resolution and low acquisition times on the order of seconds to minutes. Moreover, commercial setups are available for confocal microscope-based FCS. Compared to conventional fluorescence imaging of monolayers, the high-numerical aperture objectives employed in FCS measurements also afford higher resolution images of monolayers in comparison to air objectives. Using fluorescently labeled proteins and membrane constituents, Fluorescence Cross-Correlation Spectroscopy (FCCS) is used to quantify binding affinities of proteins to bilayers in the form of small vesicles. FCCS yielded a low micromolar *K*_*d*_ for Sar1p binding to major-minor-mix lipids (24). Here, we complement this FCCS application by quantifying the insertion areas upon binding in monolayers using the same protein and the same lipid composition.

Insertion areas were obtained from the number of protein molecules (determined by FCS) and the area increase of the monolayer upon protein insertion. The area increase was measured either directly by the film balance or, alternatively, by monitoring the dilution of a labeled lipid analog by two-color FCS.

The approach we take in this FCS application to a planar type of membrane is related to the z-scan-FCS by Benda *et al*. (15). Instead of a mechanically controlled focus position, slow evaporation of the subphase moves the monolayer through the focus (13). Since the curvature of the parabolic z-dependence of the apparent particle number and diffusion time are not used as an intrinsic calibration standard (25), small movements of the setup and the water level are not a concern for the determination of the insertion areas.

## Materials and Methods

### Lipids

Lipid monolayers were prepared either from a mixture of 1,2-dilauroyl-*sn*-glycero-3-phosphocholine (DLPC) and 1,2-dilauroyl-*sn*-glycero-3-phospho-L-serine (DLPS) at an 80:20 molar ratio or from a complex mixture of lipids (termed the ‘major-minor-mix’) that was previously established in COPII *in vitro* reconstitution experiments (26-28). It consists of 34.4 mol% of 1,2-dioleoyl-*sn*-glycero-3-phosphocholine (DOPC), 14.8 mol% of 1,2-dioleoyl-*sn*-glycero-3-phosphoethanolamine (DOPE), 3.4 mol% of 1,2-dioleoyl-*sn*-glycero-3-phosphate (DOPA), 5.4 mol% of 1,2-dioleoyl-*sn*-glycero-3-phospho-L-serine (DOPS), 5.4 mol% of L-α-phosphatidylinositol from soy (soy-PI), 1.5 mol% of L-α-phosphatidylinositol-4-phosphate from porcine brain (PI(4)P), 0.5 mol% of L-α-phosphatidylinositol-4,5-bisphosphate from porcine brain (PI(4,5)P2), 1.3 mol% of 1,2-dioleoyl-*sn*-glycero-3-(cytidine diphosphate) (CDP-DAG) and 33.3 mol% of ergosterol. All phospholipids and CDP-DAG were purchased from Avanti Polar Lipids (Alabaster, AL). Ergosterol was from Sigma-Aldrich (Schnelldorf, Germany). Lipids were dissolved in organic solvent (chloroform:methanol 2:1 (vol/vol)) and doped with 0.005 mol% of the red fluorescent lipid analog ATTO 633 DOPE (1,2-dioleoyl-*sn*-glycero-3-phosphoethanolamine labeled with ATTO 633) obtained from ATTO-TEC GmbH (Siegen, Germany).

### Protein

The GTPase Sar1p from *Saccharomyces cerevisiae* was expressed and purified as previously described (24, 27). The protein variant Sar1pS147C/C171S was prepared in the same way, but additionally labeled with Alexa Fluor 488 maleimide (Invitrogen/Thermo Fisher Scientific, Waltham, MA, USA). For labeling Sar1p, the double mutant (S147C/C171S) was used, because it can be specifically modified in position 147. This position lies on the membrane-distal surface and thus does not interfere with membrane binding. Both labeled and unlabeled Sar1p proteins were functional in a membrane binding assay (24) and with respect to GTPase enzyme activity. Protein concentrations were determined by UV absorption, using linear unmixing of the protein spectrum and the nucleotide spectrum, because Sar1p carries a GDP molecule in its active center. Protein concentrations were confirmed by a Bradford assay. Sar1p was doted with ratios of the labeled variant ranging from 0.06% to 1.16 mol%.

### Fluorescence Correlation Spectroscopy

Single-color and two-color FCS measurements were carried out on a commercial inverted confocal fluorescence microscope with a fluorescence correlation spectroscopy unit (LSM710/ConfoCor3 from Carl Zeiss, Jena, Germany). The laser powers were attenuated using an acousto-optical tunable filter to 0.5 to 27.5 μW (488 nm) and 0.25 to 0.8 μW (633 nm), all values behind the objective, for sufficient count rates while avoiding photobleaching. A long-distance C-Apochromat 40x, NA 1.1 objective with correction collar and a nominal working distance of 620 µm was used. To avoid evaporation of the immersion medium, Immersol W (Carl Zeiss, Jena, Germany) was used as the immersion medium instead of water. The dichroic mirror 488/561/633 nm was chosen as the main beam splitter, another dichroic mirror (longpass 635 nm) as the secondary beam splitter. A bandpass emission filter (505 to 540 nm) collected the signal from the green dye and a bandpass emission filter (655 to 710 nm) the red signal. Avalanche photodiodes (APDs) served as detectors. Auto- and cross-correlation and fitting procedures were performed in MATLAB (MathWorks, Natick, MA, USA). The correlation scripts were custom-written and similar to (29). The recorded count rate traces *F*(*t*) where divided into intervals with a length of one second and the autocorrelation function *G*(τ) was calculated according to Eq. 1 and fitted individually to Eq. 2 (see *e*.*g*. (30)), which represents two-dimensional diffusion in a Gaussian focus, in the range of τ = 10 µs to 0.3 s. Here τ_*D*_ and *N*_*eff*_ represent the diffusion time and effective number of particles in the focal area, respectively. The diffusion time is related to the diffusion coefficient *D* and the lateral size of the focus ω_*xy*_ as in Eq. 3.

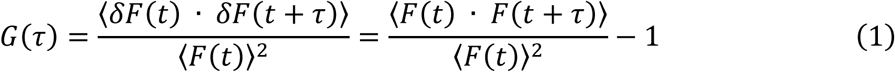

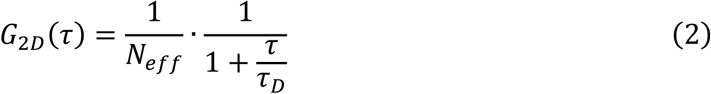

Where

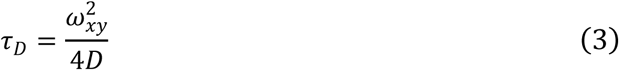

### Film Balance

The binding experiments were performed on a modified film balance from NIMA (Nima Technology Ltd., Coventry, England) (Figure 2). The Teflon trough had a maximal area of 70 cm^2^ between two moveable Delrin barriers, with a Wilhelmy plate sensor mounted to one of them. The subphase had a volume of 30 mL and consisted of a low potassium HKM-buffer (20 mM HEPES, 50 mM potassium acetate, 1.2 mM MgCl_2_, adjusted to pH = 6.8 with KOH) and 10 µM GMP-PNP at a temperature of (22 ± 0.5) °C. In addition to the thermostat connected to the trough (Alpha RA8, Lauda, Lauda-Königshofen, Germany), a second thermostat (Ecoline Staredition RE 107, Lauda, Lauda-Königshofen, Germany) was used to cool the objective. GMP-PNP acts as a non-hydrolyzable GTP-analog and keeps Sar1p in its activated state during the experiment. The trough has a circular hole in its bottom that is lined with a metal ring and sealed with a No. 1.5 coverglass (Menzel-Gl*ä*ser, Braunschweig, Germany), using a UV-cured epoxy adhesive (Loctite 3201, Henkel, Düsseldorf) to allow the long-distance microscope objective to be placed within 620 µm of the lipid film. The trough was mounted on top of the microscope to conduct FCS measurements of a film on the inverted microscope setup. To slow down evaporation, the film balance was enclosed in the Zeiss incubation housing and wet tissues were placed beside it to increase the atmospheric humidity. After the trough was cleaned with Hellmanex III, thoroughly rinsed with water and filled with buffer (verified by a pressure increase below 0.2 mN/m upon closure of the barriers), the subphase level was adjusted to the optimal height above the objective by first detecting the laser reflection at the air-water-interface with the APD detectors in LSM (laser scanning microscopy) mode. At this position the surface pressure setting was set to zero. The subphase level was raised by approximately 500 µm by adding buffer to avoid rupture of the thin water layer above the coverglass during subsequent spreading. The chloroformic lipid solution was spread using a Hamilton syringe and the solvent was allowed to evaporate for at least 30 min. Next, the barriers were used to compress the film to the desired surface pressure and its height adjusted again to the optimal position for microscopy. For FCS measurements, the lipid film was moved laterally through the microscope focus to allow acquisitions at different locations throughout the film. Each axial focal passage was recorded after raising the water level to slightly above the laser focus (as measured by the fluorescence count rate) and letting the evaporation lower it to slightly below the focus while recording FCS measurements. For measurements in the presence of protein, the film was first transferred by parallel barrier movement (*i*.*e*., at constant film area) away from the coverglass to the deeper part of the trough. Next, Sar1 was injected with a Hamilton syringe into the subphase. Subsequently, the film was moved back to the focus, again at constant film area. The subphase was stirred with two magnetic stirrers (Cimarec i Mini Stirrer, Thermo Fisher Scientific, Waltham, MA, USA) to ensure a homogeneous distribution of protein beneath the film. During the FCS measurements, the stirrers and the thermostat were temporarily turned off to avoid vibrations.

**Figure 2:**
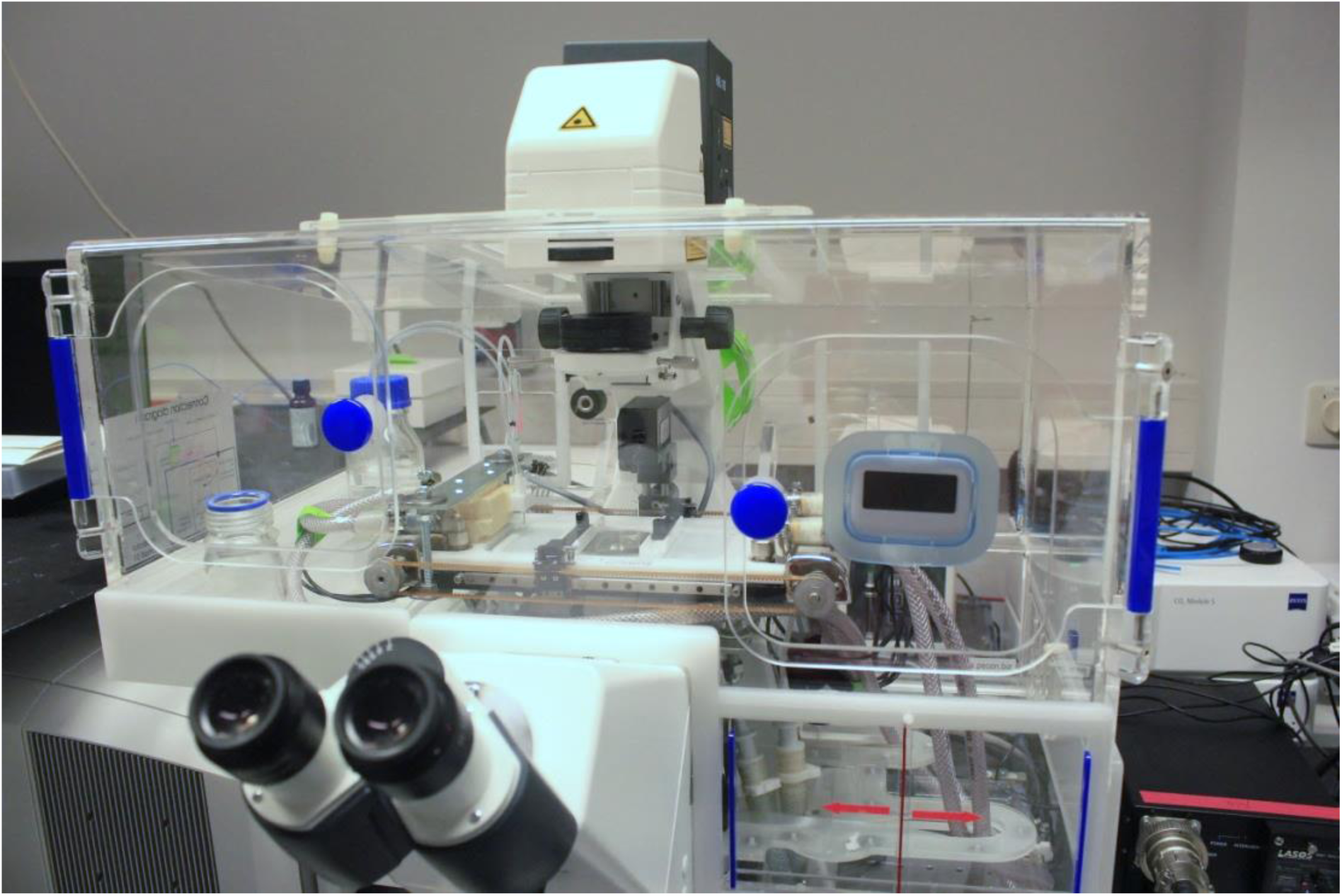
Photograph of the film balance setup on the Zeiss LSM 710/ConfoCor3 microscope, surrounded by a Zeiss incubation box. In the middle, the glued-in coverslip of the film balance with the microscope objective beneath is visible, framed on both sides by the moveable barriers of the film balance.

## Results and Discussion

### Determination of insertion areas

Sar1p protein solution was injected into the aqueous buffer below the lipid monolayer film to a final concentration of ≈ 80 nM. The resulting increase in film area at constant surface pressure was recorded by the film balance. The average insertion area per adsorbed Sar1p protein molecule was calculated by dividing the total area increase of the film by the number of protein molecules incorporated into the film.

The total number of proteins in the monolayer was obtained by extrapolating the number of protein molecules in the focal area measured by FCS to the whole film area. To this end, the absolute focal area of the confocal setup and the fraction of labeled protein need be known. To obtain an accurate measure for the focal area, raster image correlation spectroscopy (RICS, (31)) was performed (for details see Supplementary Information, Section 1). To obtain the fraction of labeled protein, the concentration of labeled protein was determined by UV/Vis-absorption, and the total protein concentration was determined in a Bradford assay.

In principle, the number of fluorescent particles in the film can be determined on a monolayer positioned axially in the middle of the focus. However, this approach proves difficult: Firstly, the monolayer needs to be positioned with high precision because any axial offset reduces the correlation amplitude and thus biases the measured particle number towards larger numbers. Secondly, it is difficult to control the height of the surface of the trough to within a few tens of nanometers by monitoring subphase evaporation and adjusting water inflow.

We therefore chose a modified type of z-scan FCS: The number of bound proteins in the focal area was determined from the fluorescence fluctuations in the monolayer during its steady movement through the focus. This movement was effected by the slow evaporation of the aqueous subphase and subsequent addition of water to start the process anew. This way, exact knowledge and control of the film position are not necessary.

After an axial focal passage had been recorded, the fluorescence trace was split into short time windows, during which the monolayer was close to stationary. Each short trace was autocorrelated and fitted with the model equation to obtain a particle number *N*_*eff*_ and diffusion time τ_*D*_ A parabolic function was fitted to the resulting parameter curves (*N*_*eff*_ vs. time and τ_*D*_ vs. time) to obtain the true parameter values, *i*.*e*., when the monolayer was in the middle of the focus.

During the axial focal passage, the extracted parameters *N*_*eff*_ and τ_*D*_ exhibited the typical parabolic form resulting from a thin fluorescent layer traveling through the laser focus (see Figure 3). This is a consequence of the form of the laser beam, which is approximately described as a focused Gaussian beam, whose squared beam width follows a parabolic dependency in the axial direction (see Supplementary Information). The minimum of the *N*_*eff*_ curve was determined by applying an unweighted least-squares parabolic fit. It represents the fluorescent particle number in the focus, which was used to calculate the insertion area.

**Figure 3:**
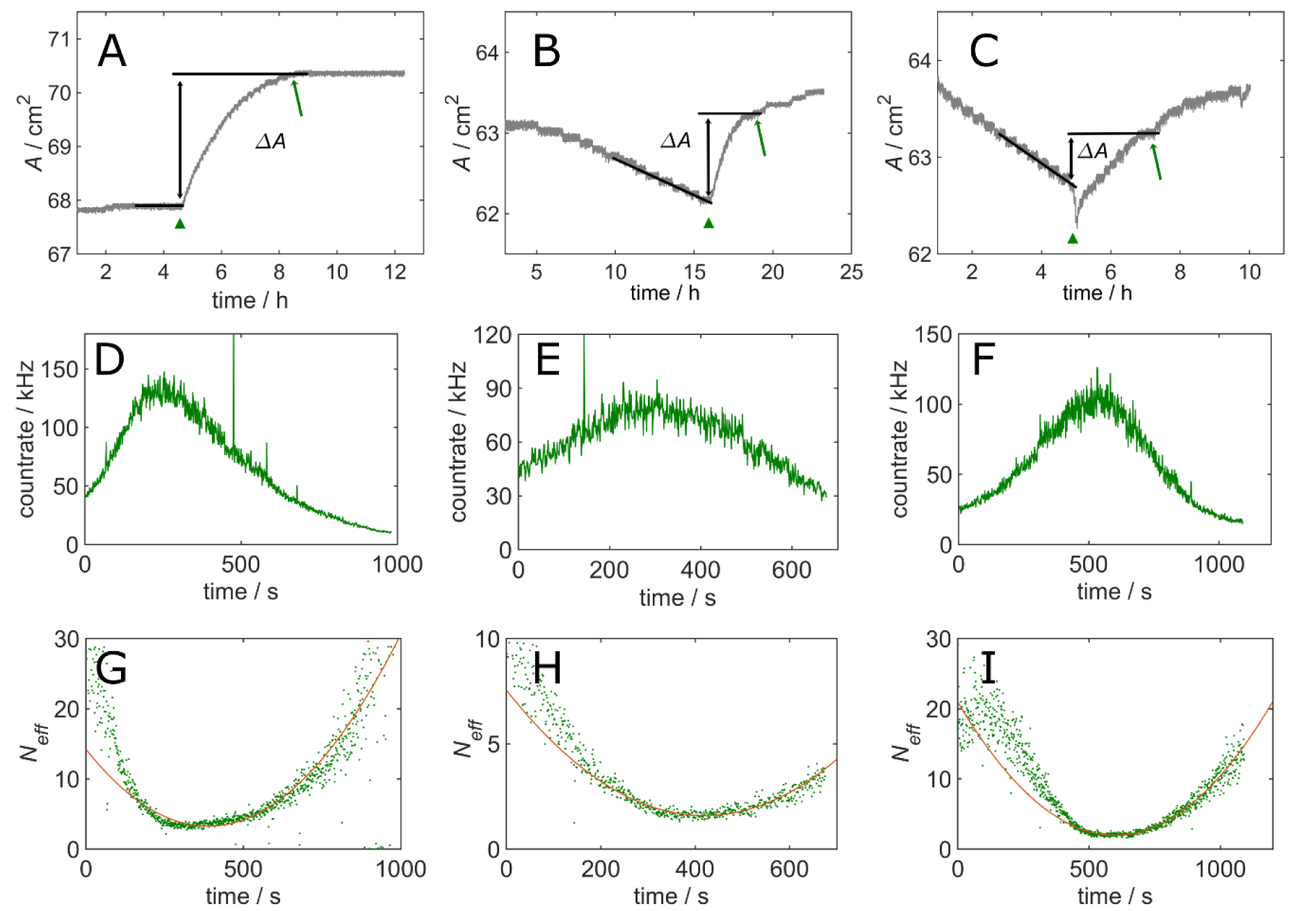
Protein insertion experiments on unlabeled monolayers from saturated lipids (DLPC/DLPS = 80:20, molar ratio). (A-C) Film balance measurements of area versus time for three independent experiments. The area increase after protein injection Δ*A* was measured as the increase in area from the time point just before injection (indicated by the green triangle) to the time at which the FCS axial focal passage was acquired (indicated by the green arrow). (D-F) Fluorescence count rate traces of the green-labeled protein during the axial focal passage on each film. (G-I) z-scan FCS: After splitting the count rate trace into 1-second subtraces, autocorrelating them and fitting with the FCS model equation, the effective particle numbers were plotted against the axial focal passage time, which serves as a proxy for the movement of the axial position of the monolayer through the focus.

Fluorescent particles diffusing in the subphase below the membrane may impact on the determination of the particle number in the focus. A detailed calculation regarding the relative influence of surface-bound and free fluorophores is provided in the Supplementary Information.

All measurements were performed on monolayers at the biologically relevant surface pressure of 30 mN/m, because the equivalence pressure at which the lipid density in monolayers and bilayers coincide is ≈ 30..35 mN/m (32, 33). Interestingly, the so-called maximum insertion pressure (MIP) of Sar1p, as determined by standard film balance measurements, for one of the lipid mixtures used (‘major-minor mix’) is also ≈ 30 mN/m. Only small amounts of protein enter the monolayer at this pressure so the corresponding area increase is very small and challenging to measure. Furthermore, at this pressure, the film is sensitive to a loss of film material, which is likely to occur at small imperfections in the trough and at barrier contact sites. Both aspects constitute a source of error in the determination of the area increase and led us to develop a method that circumvents measuring area as an extensive quantity. This is accomplished by employing FCS not only to determine the amount of inserted protein but also the concomitant reduction in lipid density. In this way, the area change due to protein insertion can be calculated even after manipulating the monolayer and possibly losing some of the monolayer by leakage underneath the barriers. The last point is important because the monolayer needed to be moved away laterally from the microscope objective to the deeper portion of the trough to permit effective stirring (see Methods section) and achieve the necessary homogeneity of protein insertion.

In each experiment, the monolayer was prepared by spreading an amount of lipid dissolved in organic solvent on the buffer surface. The solvent was allowed to evaporate and the system to reach its equilibrium surface pressure. Afterwards the film was compressed with the barriers to the final surface pressure. Due to the high pressures used in these experiments, films typically showed a steady lipid loss. The protein was injected when the area reached a slow linear decrease, while the pressure was held constant (Figure 3,Figure 4 A-C). The time required for the area to reach a maximum after injection was on the order of hours and varied between monolayer preparations. The barriers were temporarily stopped to allow FCS acquisitions without perturbations from moving the monolayer laterally. FCS measurements at different locations in the monolayer showed that the protein had distributed homogeneously.

**Figure 4:**
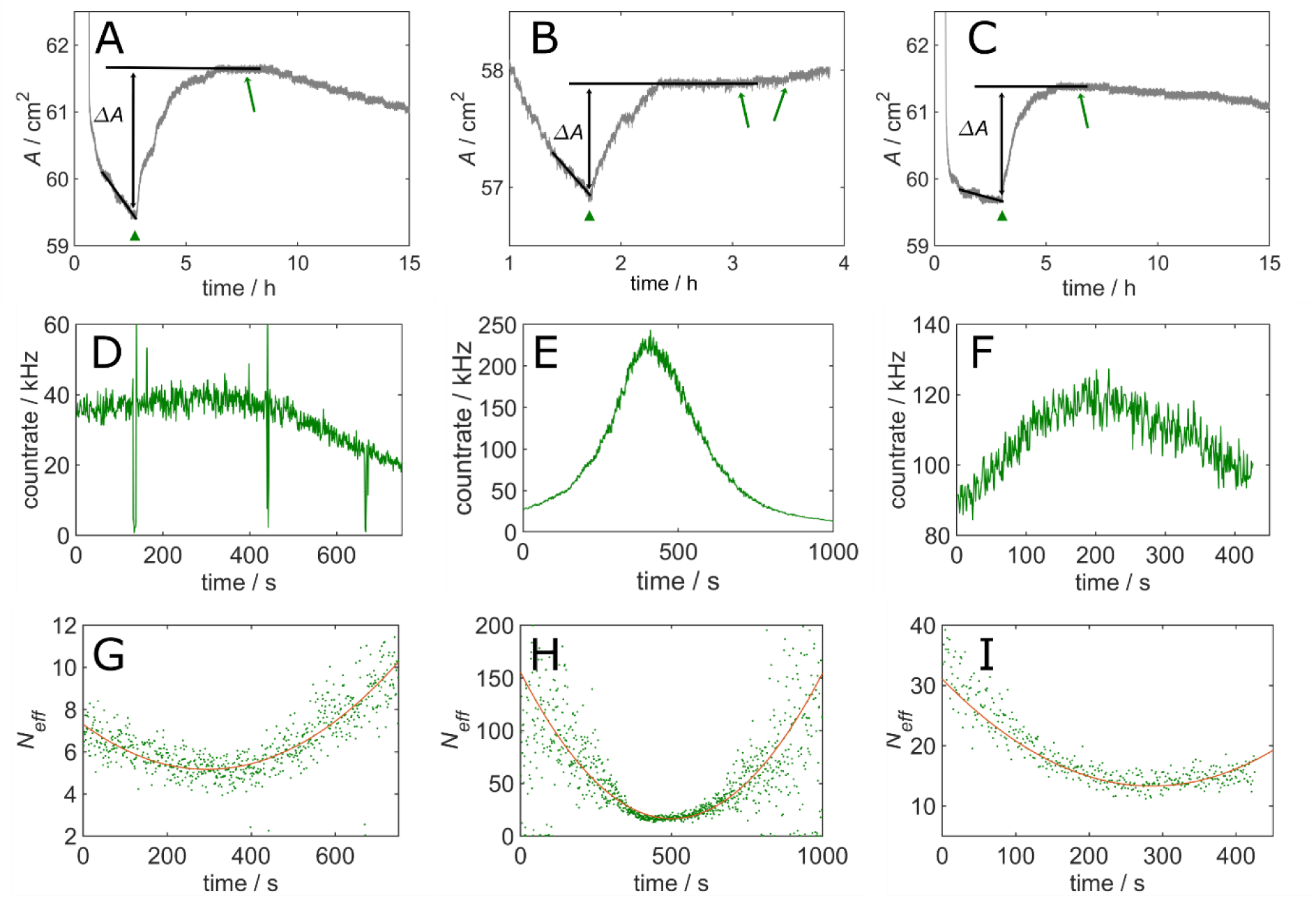
Protein insertion experiments on unlabeled ‘major-minor-mix’-monolayers. (A-C) Film balance measurements of area versus time for three independent experiments. The area increase after protein injection Δ*A* was measured as the increase in area from the time point just before injection (indicated by the green triangle) to the timepoint at which the FCS axial focal passage was acquired (indicated by the green arrow). (D-F) Fluorescence count rate traces of the green-labeled protein during the axial focal passage on each film. (G-I) z-scan FCS: After splitting the count rate trace into 1-second subtraces, autocorrelating them and fitting with the FCS model equation, the effective particle numbers were plotted against the axial focal passage time, which serves as a proxy for the movement of the axial position of the focus. Note that the different particle numbers are due to the different fractions of labeled protein used.

### Insertion into DLPC/DLPS monolayers

We started with a monolayer from a simple mixture of saturated lipids (DLPC/DLPS, 80:20 molar ratio). The insertion isotherms of three independent film preparations (Figure 3 A,B,C) all show an area increase after injection of Sar1p. Notably, the isotherm in panel A represents the ideal and rare case of a monolayer that was completely stable, both before and after protein insertion, facilitating a read-out of the area increase. The two remaining experiments show typical monolayers that diminish under constant high pressure. The area increase due to Sar1p is determined from immediately before injection to the time of the fluorescence acquisition (*i*.*e*., when the number of incorporated Sar1p molecules is measured).

The fluorescence signal of the protein that has become bound to the monolayer (Figure 3 D-F) was split into 1 second intervals and analyzed by z-scan FCS (Figure 3 G-I) as described above.

The z-scan analysis reveals a marked deviation from the expected parabolas in that they are asymmetric with respect to their minima. At least three different mechanisms may contribute to these deviations: (1) inconstancy of the evaporation rate of the subphase, leading to faster or slower movements of the monolayer through the focus; (2) influence of the air-water-interface on the shape of the detection volume (34); (3) the relatively high amount of protein in these experiments leads to a contribution of the free protein in the subphase to the correlation curves. The presence of free protein in the subphase can be seen from the reversal of the trend in the particle number *N*_eff_ towards early times in Figure 3I, when the brightness from freely diffusing protein molecules in the subphase dominates over the brightness of out-of-focus proteins adsorbed to the monolayer.

To estimate the impact of this free protein species on the FCS amplitudes, the count rates obtained during the focal passages can be used. The details of this more generally applicable theoretical framework can be found in the Supplementary Information Section 6. The theory is not limited to the binding of proteins to monolayers, but may also be employed to describe the binding of fluorescent particles (such as proteins or ligands) to the plasma membrane or to artificial bilayers such as giant unilamellar vesicles (GUV) in the presence of free fluorescent particles on either side, i.e. the interior (the cytosol) or the exterior (the extracellular space). The approach presented in the Supplementary Information is similar to the one by Smith et al. (35), but we focus here on amplitudes and particle numbers rather than particle brightness.

The three experiments shown in Figure 3 yield a mean insertion area of (11.4 ± 3.6) nm^2^. Comparing the values obtained experimentally (11.4 nm^2^) with the areas expected for the amphipathic helix alone (2.9 to 4.3 nm^2^, see Supplementary Information) and for the whole protein (13 nm^2^, see Supplementary Information) shows that not only the helix is inserted into the membrane and suggests that the whole protein becomes incorporated into DLPC/DLPS monolayers.

### Insertion into ‘major-minor-mix’ monolayers

Next, we used a more complex lipid mixture that was previously optimized for COPII-binding, the so-called ‘major-minor-mix’ (36).

Figure 4 shows film balance isotherms of three ‘major-minor-mix’-monolayers. FCS measurements were performed at the time points indicated by the arrows. The insertion area, calculated from FCS measurements on four monolayers, was (4.1 ± 2.2) nm^2^ and thus considerably smaller (by a factor of two to five) than with the DLPC/DLPS monolayers.

The experimentally determined insertion area corresponds approximately to the expected, theoretical cross-sectional area of the amphipathic helix (2.9-4.3 nm^2^, see Supplementary Information). Our results confirm the assumption that Sar1p binds membranes by embedding its amphipathic helix into the proximal monolayer. The result is also in line with a recently published cryo-TEM structure of membrane-bound Sar1p within the COPII complex (37), which finds a 90 degree bend in the amphipathic helix resulting in the lower insertion area of 2.9 nm^2^.

Examples of differences in insertion areas depending on monolayer composition and surface pressure have been previously reported (5, 7): A change from POPE to POPC monolayers doubled the area change upon adsorption of cytochrome *c* (5) and an increase of surface pressure from 14 mN/m to 18 mN/m lowered the insertion area of secretory phospholipase A_2_ from over 6 nm^2^ to nearly zero (7). The different insertion areas of Sar1p depending on the lipid composition might be due to the high sterol content of the ‘major-minor-mix’. Ergosterol (like cholesterol) reduces the fluidity of the membrane, lowering the lateral diffusion coefficient of the inserted protein and posing a greater barrier against insertion of the whole, globular protein. A driving force for a larger insertion area of Sar1p in the case of the shorter chain lipids (DLPC/DLPS) could also be the high affinity of Sar1p for the air-water-interface, as was shown in (38).

The experiments shown in Figure 3 and Figure 4 show a large uncertainty in the determination of the total area increase. To improve on this method, we introduced a second color channel to perform FCS measurements not only on the protein, but also on the lipid monolayer itself and remove the need for using the change-in-area readout of the film balance. The area increase was now calculated from the dilution of a red fluorescent lipid marker. For this purpose, 0.005 mol-% ATTO 633 DOPE was added to the lipid mixture and axial focal passage curves acquired simultaneously from the green-labeled protein and the red-labeled lipid in two detection channels. Otherwise, the experiments followed the same procedure and the number of inserted protein molecules was determined as described above.

The insertion area is calculated by dividing the area increase Δ*A* by the number of inserted proteins *N*_*P*_ according to

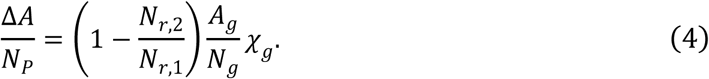

Here *A*_*g*_ denotes the area of the green focus, *N*_*g*_ and *N*_*r*_ the number of green and red particles in the focus (before insertion: *N*_*r*,1_, after insertion: *N*_*r*,2_), derived from the FCS amplitudes, *χ*_*g*_ the fraction of labeled protein. The equation is derived in the Supplementary Information Section 3.

Using this two-color FCS approach, film instabilities do not factor into the calculation, provided that the film composition remains unaltered. By measuring fluorescent protein and lipid densities in the focus (which are intensive quantities) and avoiding the use of the total area change (an extensive quantity), any film loss at the barriers (*i*.*e*., any change in system size) becomes irrelevant.

The feasibility of this approach was demonstrated on monolayers prepared from ‘major-minor-mix’ lipids (Figure 5). Z-scan FCS data was collected at multiple positions on the monolayer to extract the lipid density before and after the injection of Sar1p and the protein density after injection as accurately as possible. To do so, the monolayer was moved repeatedly relative to the fixed position of the objective on the microscope setup, by moving both barriers simultaneously. Pushing the film around causes further film loss, but the quantity does not enter the calculations, assuming that film loss is a non-selective event that does not differentially affect a subset of lipids or the protein.

**Figure 5:**
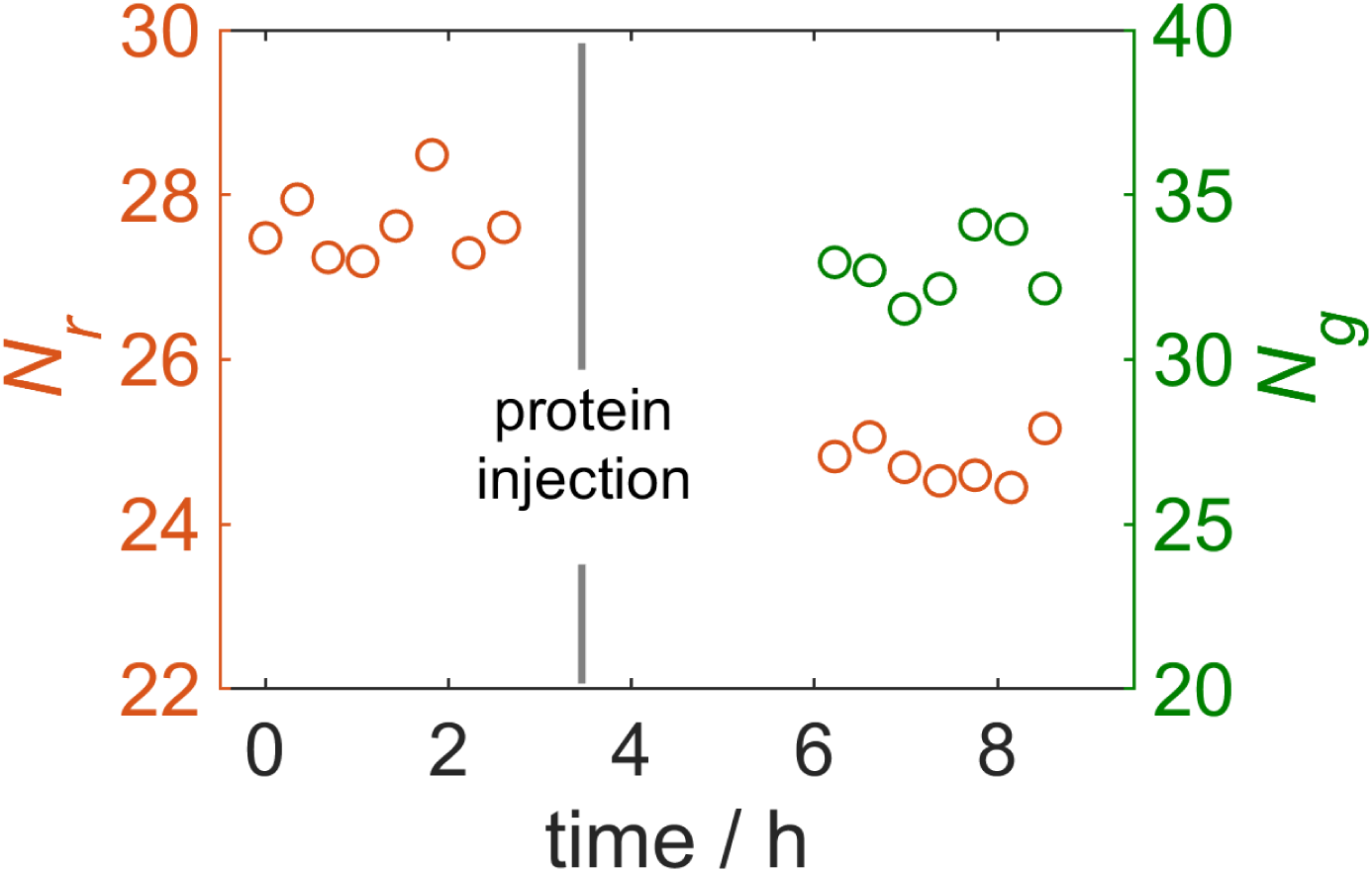
Exemplary results of green-labeled protein incorporating into red lipid-labeled ‘major-minor-mix’-monolayers. The particle numbers in both color channels are plotted against the time. Green particles are only present in the monolayer after injection. The relative change in the red particle number *N*_*r*,2_/*N*_*r*,1_ due to protein incorporation is used to calculate the relative decrease in the film area according to Eq. 4.

The resulting insertion area of Sar1p from three repetitions is (3.4 ± 0.8) nm^2^. This result is in the range of values from the single-color method (4.1 ± 2.2) nm^2^ and in agreement with the concept that only the amphipathic helix becomes embedded.

The FCS measurements additionally allowed to analyze the diffusion times of the lipid probes before and after injection of the protein. They were markedly different: Labeled lipids in the pure monolayer exhibited diffusion times of τ_*D*_ = (1.1 ± 0.1) ms. The diffusion time more than tripled upon insertion of Sar1p (τ_*D*_ = (3.4 ± 0.2) ms). This increase in diffusion time may be due to Sar1p molecules acting as moving obstacles in the monolayer (39, 40).

### Comparison with other techniques

In the literature, only few alternative approaches to determine insertion areas exist, most of which use the total area increase of the film upon insertion of the protein and are subject to the problems described above. Some authors forgo this quantity completely in favor of a purely thermodynamic calculation and fit a Gibbs isotherm to concentration-dependent increases in surface pressure to acquire the surface excess of a dissolved species and calculate an effective area from it (9). This area is not directly comparable to the lipid film area displaced by the protein and can only serve as a rough estimate (41, 42).

Other works (8, 43) used the dependency of the partition coefficient on the surface pressure to fit an exponential equation to the relative area increases at different surface pressures. This approach assumes a constant insertion area over a sufficiently large range of surface pressures, which cannot always be taken for granted (7).

When the total area increase of the film after protein injection is to be divided by the number of inserted molecules, the simplest approach is to assume that all injected molecules localize to the monolayer after equilibration (6). This assumption is justified in some cases, but since Sar1p like many other proteins readily occupies free surfaces including PTFE and the injected amount of protein is much higher than the calculated amount in the monolayer, it is not appropriate here.

Huang *et al*. (7) indirectly determined the amount of inserted protein by utilizing the protein-to-lipid ratio measured with FTIR-spectroscopy. Together with a calibration curve and the known mass of spread lipids in the film, the IR band intensities allow the calculation of the absolute mass of inserted protein. One drawback of this approach lies in the transfer of the monolayer to an ATR crystal for the IR measurement, so that the system is inevitably disturbed and any repetitions will require separate monolayer preparations. The authors find insertion areas covering a wide range of values, depending on the surface pressure, which is interpreted as the protein having different insertion modes with different penetration depths. This is analogous to Sar1p having a larger insertion area in the DLPC/DLPS film with shorter lipid tails and without ergosterol relative to the ‘major-minor mix’.

Compared to these existing methods, the FCS-based approach to determining the amount of protein bound to the membrane has several advantages:

1. The monolayer is not disturbed by the measurement and the density of bound protein is available as a function of time and position. Thus, temporal and spatial heterogeneity can be assessed to monitor for example insertion kinetics or protein-lipid interactions within phase-separated monolayers.
2. In contrast to the work of Shank-Retzlaff *et al*. (9), a series of measurements at different pressures is not required, because no presupposed functional dependency is exploited. The measurement needs to be carried out solely at the surface pressure of interest.
3. Complete binding of the protein is not a necessity in contrast to the analysis by Gerard-Egrot *et al*. (6). Therefore, the influence of varying the amount of injected protein can be studied. Furthermore, if no protein is lost (by coating of the trough or air-water-interface) the data should suffice to calculate dissociations constants and study changes in affinity as the experimental parameters are varied (e.g. surface pressure, lipid composition, temperature, ion strength). However, in most cases the assumption is not justified (see for example (44, 45)).
4. In addition, the dynamic interaction of the proteins with the monolayer can be analyzed with respect to diffusion coefficients and photophysics.

Additional advantages arise when the measurement of the area increase with the film balance is replaced by a measurement of the relative change in the lipid dye density by FCS:

1. Film instabilities that distort the true area increase are no longer a concern, because only the protein and lipid densities in the focal area contribute to the calculation.
2. Insertion areas could also be determined in restricted geometries. Protein-monolayer interactions including protein insertions areas could be studied locally in different phases of phase-separated systems.
3. The method could be extended to flat or nearly-flat bilayer membranes, for example in giant unilamellar vesicles (GUVs) or supported bilayers to determine protein insertion areas also in bilayer membranes.

The Langmuir film balance method is typically used to characterize a protein’s propensity for membrane insertion by determining its so-called maximum-insertion pressure (MIP) (46). This is the pressure at which the macroscopic film balance measurement reveals no further increase in monolayer pressure upon protein injection at constant area. By virtue of its single-molecule sensitivity, FCS also paves the way for elucidating protein-lipid interactions at and potentially slightly beyond the MIP. For Sar1p, protein-lipid interactions at the MIP in ‘major-minor-mix’ monolayers are of particular interest, because the MIP coincides with the physiologically relevant monolayer-bilayer equivalence pressure (38) and its ability to insert at this pressure may be linked to its membrane bending activity on bilayers. It remains to be investigated if this is a general mechanism used by amphipathic helix proteins that function in membrane remodeling during intracellular trafficking.

## Conclusion

We have introduced a method to measure insertion areas of membrane proteins in lipid monolayers that employs two-color FCS and does not rely on macroscopic area measurements. The lipid film is probed locally. Further assumptions that were required in previous works, for instance regarding the fraction of bound protein, are no longer necessary.

We have applied this method to the insertion of the protein Sar1p into lipid monolayers and were able to distinguish two putative insertion mechanisms depending on the lipid composition. The data is consistent with only the amphipathic helix of Sar1 inserting into the less fluid ‘major-minor-mix’ monolayers and the whole protein incorporating into monolayers made from shorter, saturated lipids.

The theoretical framework presented in this work also paves the ground towards analyzing insertion areas in bilayer membranes. Moreover, theoretical considerations for accounting for the contributions from membrane-bound and freely diffusing fluorescent particles to an FCS amplitude are included.

## Author Contributions

K.B. designed research, A.M. and J.A. established the setup, J.A. prepared the samples and performed measurements, J.E. and C.S. performed additional experiments, J.A. and J.E. analyzed the data, A.S. calculated the theoretical insertion areas, J.E. and K.B. wrote the manuscript.

## Acknowledgements

The BMBF (03Z2HN22), Land Sachsen-Anhalt/ERDF (124112001 and 1241090001) and the DFG (FOR1145) are acknowledged for funding.

## Supporting Citations

References (2, 24, 30, 31, 36, 37, 47-52) appear in the Supporting Material.

